# Weakly Supervised Vector Quantization for Whole Slide Image Classification

**DOI:** 10.1101/2024.08.31.610626

**Authors:** Dawei Shen, Yao-zhong Zhang, Seiya Imoto

## Abstract

Whole Slide Images (WSIs) are gigapixel, high-resolution digital scans of microscope slides, providing detailed tissue profiles for pathological analysis. Due to their gigapixel size and lack of detailed annotations, Multiple Instance Learning (MIL) becomes the primary technique for WSI analysis. However, current MIL methods for WSIs directly use embeddings extracted by a pretrained vision encoder, which are not task-specific and often exhibit high variability. To address this, we introduce a novel method, VQ-MIL, which maps the embeddings to a discrete space using weakly supervised vector quantization to refine the embeddings and reduce the variability. Additionally, the discrete embeddings from our methods provides clearer visualizations compared to other methods. Our experiments show that VQ-MIL achieves state-of-the-art classification results on two benchmark datasets. The source code is available at https://github.com/aCoalBall/VQMIL.

## I. Introduction

Whole Slide Images (WSIs) are high-resolution digital scans of entire microscope slides, widely used in pathology. [1] In recent years, deep learning methods for analyzing these WSIs have advanced rapidly, achieving performance on par with or even exceeding that of pathologists. This progress enables scalable and thorough examination of tissue samples in clinical practice.

One of the key applications of deep learning methods for WSI analysis is WSI tumor classification, which focuses on identifying the presence of tumor tissues within WSIs. This classification task is well-suited for Multiple Instance Learning (MIL)[2, 3], a form of weakly supervised machine learning that organizes data into labeled bags, each containing multiple instances. This method is advantageous for WSI tumor classification for two reasons. First, the gigapixel size of WSIs makes it infeasible for current hardware and algorithms to process them directly. This necessitates dividing each WSI into numerous smaller image tiles and compressing each tile into a small embedding vector. Second, detailed annotations for WSIs are scarce, as providing pixel-level annotations for each slide is costly and time-consuming for pathologists.[4, 5] Hence, training usually utilizes slide-level labels rather than detailed instance labels.

The current MIL process for WSIs begins with a pretrained vision encoder, such as ResNet [6] or vision transformer [7], which compresses each image tile into an embedding vector. These vectors represent the instances in the MIL framework. A MIL model is then trained on these embedding vectors to perform slide classification. This process efficiently addresses the challenges of size and annotation scarcity inherent in WSI tumor classification. A key factor of this process is the quality of the embedding vectors.

However, the embedding vectors obtained from pretrained vision encoders have two drawbacks. The first issue is that the embeddings are not task-specific. The pretraining frameworks are designed to learn general features from a large and diverse dataset, rather than focusing on a specific task like tumor classification. The second issue is the high variability of the continuous embedding vectors. Continuous embeddings from the vision encoder introduce extraneous variations, such as subtle differences in texture or appearance between tiles, impairing the MIL model’s ability to perform accurate classification.

To address these two issues, we propose VQ-MIL, which incorporates a weakly supervised vector quantization module between the vision encoder and the MIL model. The module includes a learnable codebook with a fixed number of codewords. During training, embedding vectors from the pretrained vision encoder are refined by an additional encoder to highlight the task-specific information, then these embeddings are mapped to the nearest codewords. These discretized codewords, rather than the original embedding vectors, are passed to the MIL model. This process can be treated as a supervised clustering process that reduces variability of the embeddings and thereby enhancing the MIL model’s classification performance. In III-B2, we further explain the effectness of our method using the information bottleneck principle [8]. Additionally, the trained codebook has fixed values, providing clearer and more accurate visualizations compared to continuous embedding vectors used in previous methods[9, 10].

Furthermore, we argue that another bottleneck for WSI classification is the lack of slide-level labels. To address this, we randomly select image tiles from tumor and normal slides to create positive and negative pseudo-bags as a simple data augmentation method.

We evaluated our method on two representative datasets, Camelyon16 [11] and TCGA lung cancer dataset, and the experimental results show that our model achieves state-of-the-art slide-level performance compared with the previous methods.

To our knowledge, this is the first use of vector quantization in weakly supervised classification. It demonstrates that vector quantization can refine the embedding space for WSI classification, improving diagnostic accuracy.

## II. Related Works

### A. MIL Methods for WSI Classification

To address the challenges of gigapixel size and the lack of precise annotations in WSIs, current deep learning methods typically employ a two-stage approach that consists of a vision encoder followed by a MIL model. In this process, a WSI is initially divided into many smaller image tiles. A pretrained and frozen vision encoder is then used to convert each of these tiles into one-dimensional embedding vectors. There are two main types of pretrained vision encoders. The first type is a general-purpose vision encoder pretrained on datasets like ImageNet. However, there is a significant domain gap between ImageNet images and pathological images. To address this issue, some research [12, 13, 14, 15] has focused on using a large number of unlabeled WSIs to pretrain vision encoders within self-supervised learning frameworks such as SimCLR [16]or DINO [17, 18].

After extracting the embedding vectors, all vectors from each WSI are collected to train a MIL model that acts as a slide-level classifier. There has been significant work in developing efficient MIL models for WSI classification. AB-MIL [9] introduces a gated-attention mechanism [19] to aggregate features using attention scores. CLAM [10] is a variant of AB-MIL that clusters instances with the top-k highest and lowest attention scores. DS-MIL [12] combines embedding vectors from multiple scales and selects the most representative instance as a query to identify other instances. TransMIL [20] utilizes Transformer[21] architecture to consider the relationships between instances in a slide. DTFD-MIL[22] resamples instances within a slide to increase the number of available labels. MambaMIL[23] utilizes the Selective Scan Space State Sequential Model (Mamba)[24] architecture to process long sequences of instances. All of these methods demonstrate reliable performance in slide-level classification.

Recently, there have been some impressive efforts[25, 26, 27] to attempt to finetune the vision encoders to address the issue that the embeddings generated by the vision encoder are not task-specific. Considering the powerful capabilities of vision encoders, this approach has great potential. However, due to the enormous size of WSIs, it is not feasible to finetune the vision encoder using entire WSIs. To mitigate this issue, some studies[26] finetune the model by assigning the slide-level label to each instance within the slide, while some other studies[25, 27] reduce the input size by sampling instances using methods like K-Means[28] or deep variational information bottleneck[29]. However, these methods inevitably result in incomplete or incorrect data, which can impair the model’s performance comparing with fully-supervised finetuning.

### B. Vector Quantization

Vector quantization is a technique for compressing data by mapping vectors in high-dimensional space to a finite set of prototype vectors or codebook, effectively reducing the data’s dimensionality while preserving its essential features.

Vector Quantized Variational Autoencoder (VQ-VAE)[30] is a variant of variational autoencoder (VAE)[31] that utilizes vector quantization. It encodes input data into discrete latent space by mapping continuous features to the nearest codeword in a learnable codebook. Then this discrete representation is used to generate high-quality reconstructions. VQ-VAE is particularly useful for tasks like image generation and speech synthesis, where capturing discrete underlying structures can improve the model’s performance and interpretability.

In our work, unlike VQ-VAE which employing vector quantization to reconstruct input data, we employ it to improve the task-specificity of embeddings, mapping them to a discrete space that aligns better with weakly supervised classification. By integrating this into the MIL pipeline, we aim to improve classification accuracy and instance-level distinguishability.

## III. Methods

### A. Formulation of MIL Methods for WSI

To formalize the process of WSI deep learning analysis, denote the frozen vision encoder as *V*_*e*_. For an input WSI X = {*x*_1_, *x*_2_, …, *x*_*k*_}, where *k* is the total number of tiles and *x*_*i*_ is the *i*^*th*^ tile that cropped from the WSI. The set of embedding vectors F = {**f**_1_, **f**_2_, …, **f**_*k*_} is generated by *V*_*e*_ that **f**_*i*_ = ∈*V*_*e*_(*x*_*i*_), **f**_*i*_ ℝ^1*×N*^, *N* is the length of **f**_*i*_. F is regarded as the instances of slide X in the settings of MIL. Then the slide label Y of X, is determined by the latent instance labels {*y*_1_, *y*_2_, …, *y*_*k*_}:

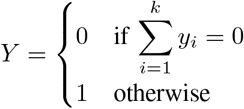

Since the labels of instances are unknown, usually a MIL model *A* is applied on F to produce slide-level prediction 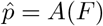.For gated-attention[19] based MIL models like AB-MIL[9], a learnable attention score *a*_*i*_ is calculated for each instance **f**_*i*_ in the following way:

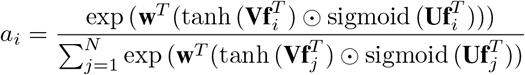

where ⊙ is the element-wise multiplication operation, **w, U** and **V** are learnable parameters. **U, V**∈ ℝ^*M ×N*^, **w**∈ ℝ^*M ×*1^, that is, **U** and **V** linearly map the input from dimension *N* to *M* .

All instances in a slide are then aggregated as a slide-level embedding and a linear head is used to do the prediction. the model could be described as :

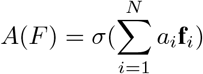

where *σ* is the linear classifier head.

### B. VQ-MIL

In this section we introduce our model VQ-MIL, especially the vector quantization module in it and how it maps the embedding vectors to a discrete and task-specific space. The overview workflow of our method is shown in Figure 1. We split our workflow into three modules, the vision encoding module, the vector quantization module and the MIL module. In the vision encoding module, the input WSI *X* is first divided into N tiles. A vision encoder *V*_*e*_ independently encodes these N tiles into N embedding vectors. Note that these N embedding vectors are general and not task-specific. They are then passed to the vector quantization module, where a learnable encoder *Z*_*e*_ refines them. As *Z*_*e*_ is learnable, it non-linearly transforms these input embedding vectors, highlighting the task-specific information. Then there is a learnable codebook *Q* that contains K discretized codewords. All these N embedding vectors are respectively mapped to the nearest codeword. Then it is the mapped codewords instead of the embedding vectors are passed to the MIL module. Here we simply apply the classic AB-MIL as the MIL model *A*, in which an attention score is calculated for each codeword and all codewords are aggregated as a slide embedding by the weight of attention scores. A linear layer is applied on the slide embedding to do slide prediction.

**Fig. 1:**
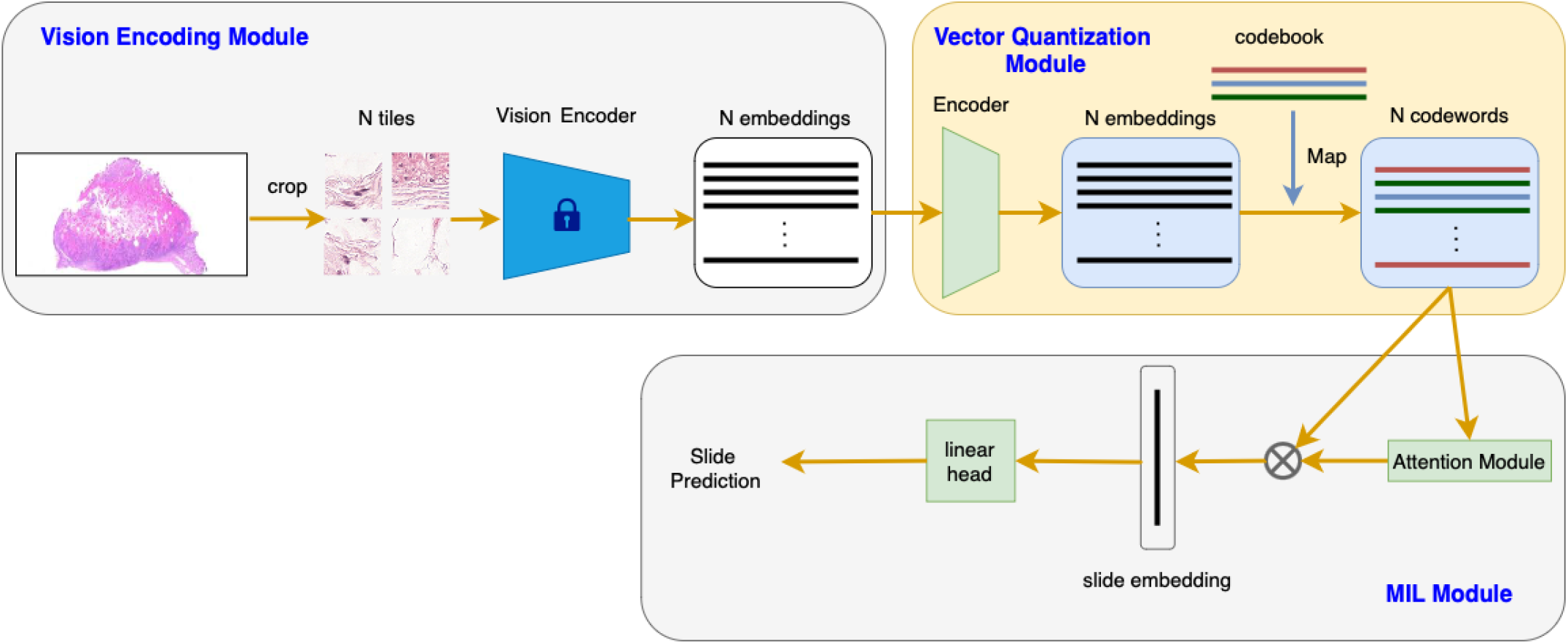
The workflow of VQ-MIL, which have three modules. The vision encoding module converts a WSI to embedding vectors. The vector quantization module mapped embedding vectors to discrete and task-specific codewords. The MIL module aggregates all codewords to make the prediction.

#### 1) Vector Quantization Module

Inspired by VQ-VAE, we tried to apply vector quantization in MIL to map the continuous embedding vectors to a discrete space. Here, to apply the vector quantization method on WSI classification problems, we add an additional vector quantization module after the conventional vision encoding module and before the MIL module.

In the vector quantization module, a codebook is defined as *Q*∈ *R*^*K×D*^, where *K* is the size of the codebook and *D* is the dimension of each codewords. For a WSI X, the generated embedding vectors F are passed through the vector quantization module, where an additional encoder *Z*_*e*_ further encodes the embeddings as E = {**e**_1_, **e**_2_, …, **e**_*k*_}, where **e**_*i*_ = *Z*_*e*_(**f**_*i*_). Then the set of codewords C ={**c**_1_, **c**_2_, …, **c**_*k*_} is calculated in the way that for each **e**_*i*_, **c**_*i*_ is the nearest codeword from it:

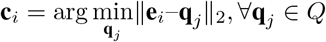

Then it is the calculated set of code words C instead of F is passed through the MIL model *A* to do the prediction 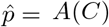.C is in a discrete space since it is a set of codewords from the codebook.

Here we face the problem that the vector quantization process is not differentiable. We apply the same method as in VQ-VAE[30] that directly copying the gradient from *C* to *E*. Thus the actual expression of the workflow is:

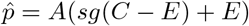

where *E* = *Z*_*e*_(*F*). *sg*(*x*) = *x* during forward propagation and *sg*(*x*) = 0 during back propagation.

Besides the classification loss, we also add two vector quantization loss similar to which in VQ-VAE that updates both the outputs of the encoder and the codewords to make them closer.

Therefore the loss function is:

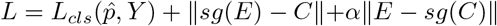

where *Y* is the slide label and *α* is a coefficient. The first term is the slide-level cross-entropy loss, the second and the third term are vector quantization loss. During the training, the parameters of the vector quantization module and the MIL module are updated simultaneously.

Note that compared with VQ-VAE[30], our VQ-MIL replaces the decoder with the MIL model and replaces the reconstruction loss from the decoder with the weakly supervised slide-level classification loss from the MIL model.

Algotithm1 is the pseudo code of the workflow.

##### Algorithm 1 pseudo-code of VQ-MIL

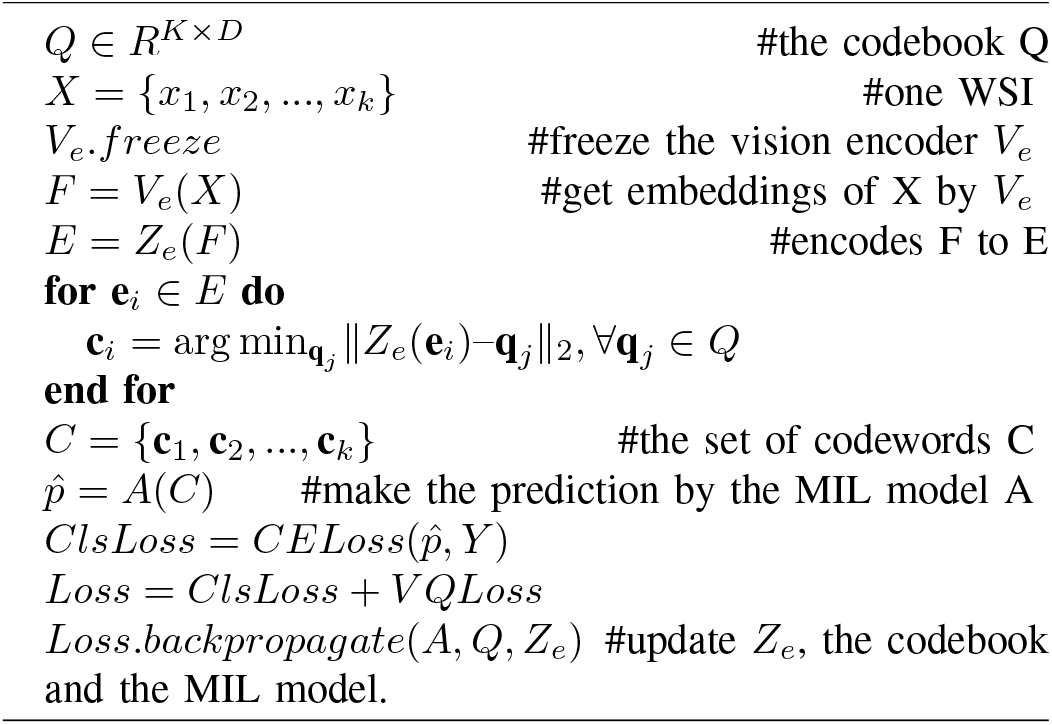

#### 2) Vector Quantization and Information Bottleneck Principle

Here, we use the information bottleneck principle to explain the effectiveness of vector quantization. In supervised learning, input variable *X* is often complex and high-dimensional while label variable *Y* is simple and low-dimensional, meaning much of the information in *X* is irrelevant to *Y* [8]. According to Tishby et al. (2015), a key goal of supervised learning is to capture and efficiently represent the features in *X* that are relevant to *Y* . Therefore, the information bottleneck principle can be applied to analyze these models. The information bottleneck principle is a theoretical framework trying to find the embedding 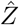 that minimizes the following *B*:

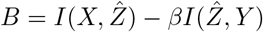

where *X* is the input variable, *Y* is the label variable, *β* is an coefficient and the function *I* is the mutual information which mathematically measures the amount of information one variable contains about another.

In other words, the information bottleneck principle requires the embedding 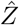 to retain more information relevant to *Y*, but reduce irrelevant information.

In VQ-MIL, the vector quantization module acts as a weakly supervised clustering process. Guided by slide-level labels, embedding vectors are non-linearly clustered around codewords with only the codewords proceeding to the next steps. As a result, 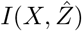 is implicitly reduced, while 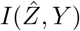 is maintained by the weakly supervised classification loss.

To further illustrate this, we conduct a comparison experiment by replacing the vector quantization module with K-Means (See IV-G3). The results show that vector quantization module effectively reduces irrelevant details, while unsupervised clustering like K-Means only converts general features from continuous to discrete.

### C. Random Pseudo Bags

One challenge in training an efficient model for WSI classification is the scarcity of labels. Given the rich and varied content within WSIs, having only one label per slide provides too little information for the MIL model to effectively exploit useful features. For instance, in the Camelyon16 dataset, which contains WSIs of lymph node sections, there are 270 slides for training and 400 slides in total. Usually, thousands of image tiles can be cropped from a single slide, resulting in only 270 labels for hundreds of thousands of tiles in Camelyon16. This lack of labels means that, in gated-attention[19] based MIL models, only a subset of positive tiles may be identified from the slide. This limitation hampers the MIL model’s ability to fully learn and distinguish the features of tumor tissues.

Inspired by previous methods[22, 32, 33], we randomly select *M* instances each time from each slide type (e.g., tumor and normal) to create a large number of pseudo-bags. This approach helps our model explore as many potential instances as possible.

## IV. Experiments

### A. Datasets and Preprocessing

We report our results on two commonly used WSI datasets: Camelyon16[11] and the TCGA lung cancer dataset.

#### Camelyon16

The Camelyon16 dataset is a benchmark in computational pathology, used for developing algorithms to detect breast cancer metastases in lymph node tissue sections. It includes 400 whole-slide images with detailed annotations provided by expert pathologists. The dataset is divided into 270 training slides from two medical centers and 130 test slides from another center. This dataset was created for the Camelyon16 Grand Challenge to evaluate the performance of automated systems in improving cancer diagnosis. Camelyon16 also provides pixel-level annotation for each slides for detailed evaluation.

#### TCGA lung cancer

This dataset includes WSIs for two major subtypes of lung cancer: lung adenocarcinoma (LUAD) and lung squamous cell carcinoma (LUSC). It contains a total of 1,054 slides, with 531 from LUAD and 513 from LUSC. The dataset can be downloaded from the National Cancer Institute Data Portal.

For data preprocessing, we followed the CLAM[10] library guidelines, using the OTSU[34] algorithm for segmentation and tile cropping. The tile size was set to 224 × 224 at 20× magnification.

### B. Evaluation Metrics

For Camelyon16, the provided training set was split into training and validation sets with a ratio of 0.9:0.1. We conducted five cross-validation experiments for a fair comparison. For the TCGA lung cancer dataset, we randomly split the dataset into training, validation, and test sets with ratios of 0.65, 0.1, and 0.25, respectively. We conducted three cross-validation experiments for this dataset. We report the mean and standard deviation of accuracy and AUC of our method alongside all baselines on both datasets.

### C. Implementation Details

For a fair comparison, we use an ImageNet-pretrained ResNet50[6] as the frozen vision encoder to generate embedding vectors for our method and all baselines. It encodes image tiles of size 224 × 224 into embedding vectors of size 1024.

For all baselines, we used the same settings as described in their respective papers and source code.

For our method, we use a 3-layer MLP as the encoder *Z*_*e*_, which projects the embedding vectors from size 1024 to size 512. We set the number of codewords in the codebook to 32, resulting in a codebook size of 32 × 512. The optimizer used is AdamW[35] with a learning rate of 0.0001. The batch size is set to 1, meaning only one slide is processed per iteration. Due to the different sizes of the datasets, we trained our model for 400 epochs on Camelyon16 and 100 epochs on the TCGA lung cancer dataset. We use pseudo-bags of size 256 as an additional data augmentation technique.

### D. Slide-Level Performance

We compare our models with the following baselines: (1) gated-attention[19] based methods, including the classic AB-MIL[9] and its two variants, CLAM[10] and DTFD-MIL[22]; the transformer-based method, TransMIL[20]; and (3) the latest Mamba-bsed method MambaMIL[23]. II-A provides brief introductions to these methods.

Table I summarizes the slide-level performance of the baseline models and our method on Camelyon16. The results show that our method improved accuracy by 5.11% and AUC by 8.66% compared to the classic AB-MIL. Our method also outperforms CLAM and TransMIL. Compared to DTFD-MIL,our method achieved similar accuracy but improved AUC by 1.01%.

**TABLE I:** Comparison of MIL workflows on Camelyon16. The standard deviations are in parentheses. The best scores are in bold.

| Model | Accuracy | AUC |
| --- | --- | --- |
| AB-MIL | 0.8326 ($\pm 0.043$) | 0.8382 ($\pm 0.021$) |
| CLAM | 0.8465 ($\pm 0.029$) | 0.8766 ($\pm 0.030$) |
| TransMIL | 0.8775 ($\pm 0.014$) | 0.8883 ($\pm 0.017$) |
| DTFD-MIL | **0.8837** ($\pm 0.010$) | 0.9147 ($\pm 0.011$) |
| MambaMIL | 0.8605 ($\pm 0.024$) | 0.9106 ($\pm 0.007$) |
| Ours | **0.8837** ($\pm 0.022$) | **0.9248** ($\pm 0.014$) |

For TCGA lung cancer dataset, table II summarizes the slide-level results. Our method achieved the best performance, with an improvement in accuracy of 4.11% and AUC of 5.36% compared to the classic AB-MIL.

**TABLE II:**
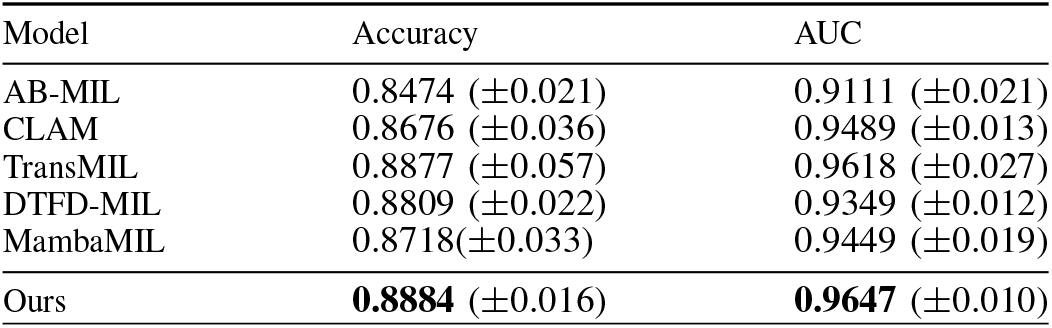
Comparison of MIL workflows on TCGA Lung Cancer Dataset. The standard deviations are in parentheses. The best scores are in bold.

### E. Distinguishability between the Tumor Instances and Normal Instances

We further tested the ability of the vector quantization module to reduce variability and distinguish between tumor and normal instances. We calculated the proportion of tumor instances corresponding to the 32 codewords of VQ-MIL trained on Camelyon16 and examined the attention scores associated with these codewords. The results are shown in Figure 2. We can see that different codewords have vastly different proportions of tumor instances. Some codewords contain more than 90% tumor instances, while many others consist almost entirely of normal instances. This indicates that codewords can effectively provide the task-specific information like distinguishing between tumor and normal instances. Additionally, there is a strong correlation between the proportion of tumor instances corresponding to a codeword and its attention score. Codewords with a high proportion of tumor instances correspond to high attention scores, whereas those with a low proportion correspond to low attention scores. This suggests that the model could effectively notice the codewords with higher tumor proportions for classification prediction.

**Fig. 2:**
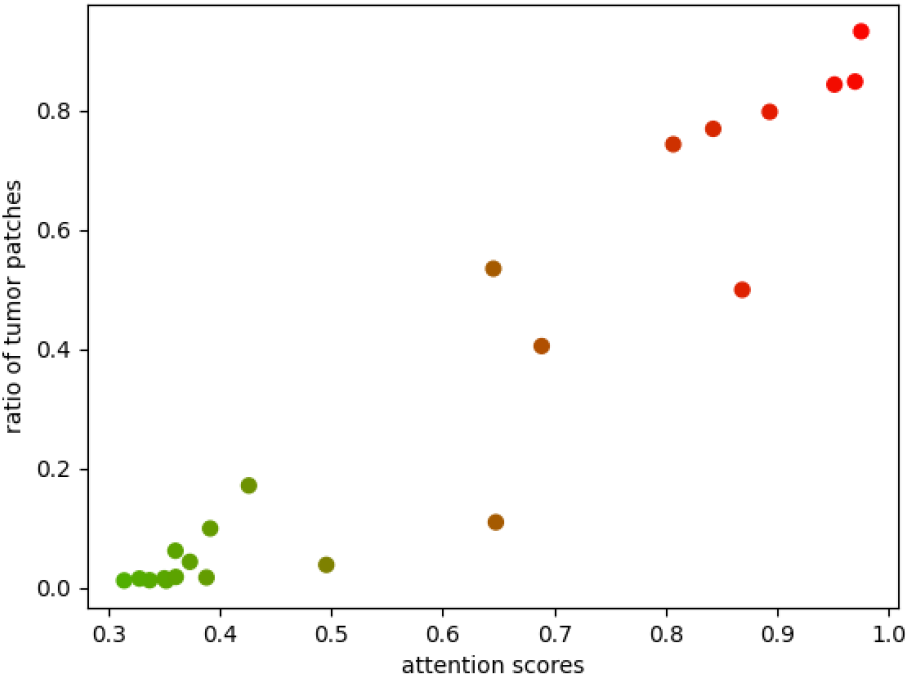
The relationship between the proportion of tumor instances of each codeword and the attention scores. Codewords with higher attention scores are red, and codewords with lower attention scores are green.

Figure 3a and 3b show the t-SNE plots of instance embedding vectors from a WSI in Camelyon16 and the slide embedding vectors obtained from all WSIs in the training set, respectively. In 3a, we observe that the instance embedding vectors generated by the vision encoder contain significant noise, with tumor instances buried among normal instances, making them difficult to distinguish. However, after being discretely mapped to codewords by the vector quantization module, some tumor instances are separated from the normal instances. This separation allows the MIL model to notice these distinct tumor instances, leading to stronger distinguishability in the slide embedding vectors, as reflected in 3b.

**Fig. 3:**
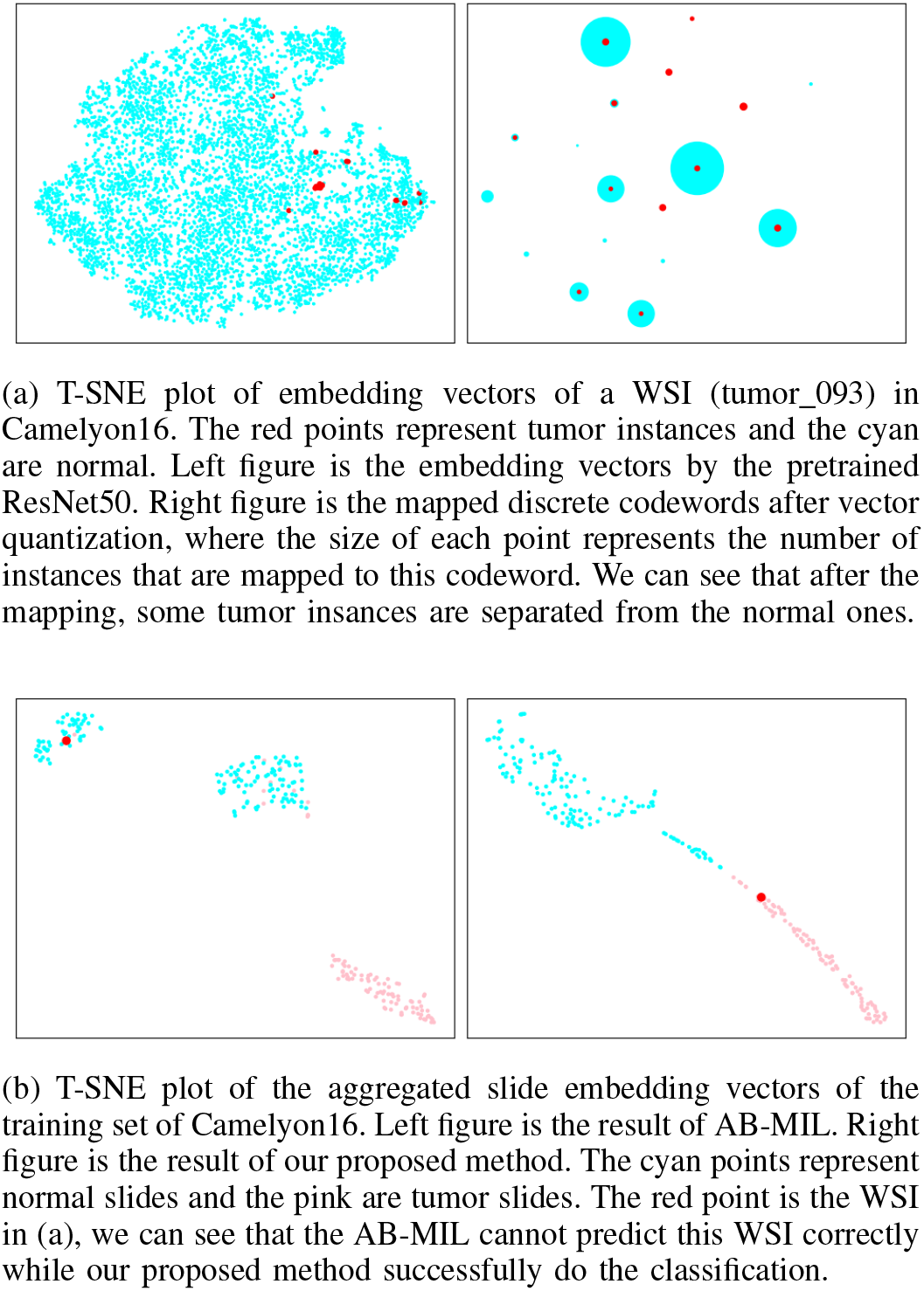
The t-SNE plot of embedding vectors in Camelyon16.

### F. Instance-Level Annotation and Visualization

In this section, we demonstrate that our method provides better visualization for instance-level annotation of WSIs compared to previous models. Currently, drawing heatmaps of attention scores using gated-attention[19] based MIL models is the common method for instance-level visualization in WSI classification. Compared to the previous gated-attention[19] based models, our method has an advantage for visualization. In our model, the attention scores derived from the code-words are fixed. In most models, attention scores has to be normalized in the model so that the sum of the attention scores for all instances in a slide is 1. This process can be detrimental to visualization. For example, higher attention scores usually should indicate a higher likelihood of being a tumor instance. However, due to normalization, there is no clear threshold for this likelihood, even normal slides may have many instances with high attention scores. In our model, although normalization is applied during prediction, the codebook is fixed after training. Thus we can use this codebook as a bag with *K* instances (codewords) and input it into the MIL model to obtain fixed attention scores. This approach results in better visualization outcomes. Figure 4 illustrates the difference in visualization performance.

**Fig. 4:**
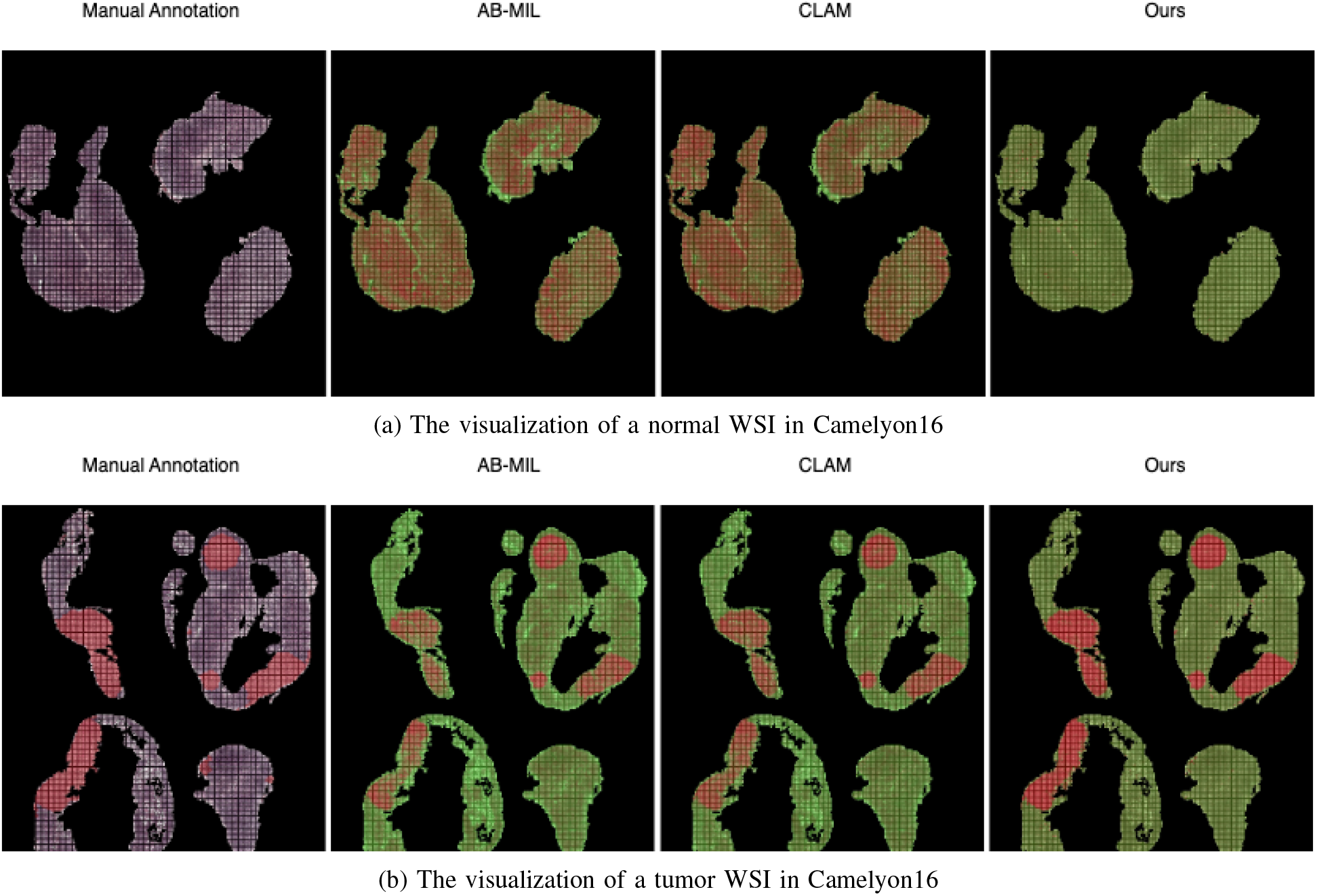
Two visualization examples in Camelyon16. The left-most column is the provided ground truth annotation. The second column is the heatmap from the classic AB-MIL. The thrid column is the heatmap from CLAM. The right-most column is the heatmap from our method.

### G. Ablation Study

#### 1) Contribution of each component

We tested the three components of our VQ-MIL:the encoder *Z*_*e*_, the vector quantization process, and the data augmentation through pseudo bags.

Table III summarize the contribution of each component on Camelyon16. In the table, the first row represents the basic AB-MIL model, which shows the lowest performance. The second row features AB-MIL with an additional encoder *Z*_*e*_. The encoder *Z*_*e*_ encodes the embedding vectors to be more task-specific, although there is no vector quantization process to map the vectors to a discrete space. This addition improved the AUC by 2.77% and accuracy by 2.96% compared to the basic AB-MIL, indicating the importance of converting embedding vectors to be more task-specific. The third row represents VQ-MIL without additional data augmentation. It further improved the AUC by nearly 3% and accuracy by 1.24%, suggesting that a discrete embedding space is beneficial. The last row is the full version of our method, which achieves the best performance, highlighting the contribution of data augmentation through pseudo-bags. In conclusion, we find that each component significantly contributes to the overall performance.

**TABLE III:** Contribution of different components of VQ-MIL on Camelyon16. Best scores are in bold.

|  | Accuracy | AUC |
| --- | --- | --- |
| AB-MIL | 0.8326 | 0.8382 |
| Encoder+AB-MIL | 0.8682 | 0.8659 |
| VQ+AB-MIL | 0.8806 | 0.8955 |
| VQ+AB-MIL+pseudo-bags (ours) | **0.8837** | **0.9248** |

#### 2) Impact of different codebook size

Table IV shows the slide-level performance on the Camelyon16 dataset concerning different codebook sizes. For binary classification tasks, it might seem intuitive to set the codebook size to two; however, the performance is unsatisfactory because the encoder of the vector quantization module struggles to accurately cluster all instances into just two codewords. Increasing the size of the codebook improves performance. We also find that there is almost no difference in performance when the codebook size exceeds 16.

**TABLE IV:** Different size of the codebook on Camelyon16.

| size | Accuracy | AUC |
| --- | --- | --- |
| 2 | 0.8465 | 0.8432 |
| 8 | 0.8202 | 0.7842 |
| 16 | 0.8620 | 0.9080 |
| 32 | 0.8806 | 0.8955 |
| 64 | 0.8806 | 0.8885 |
| 128 | 0.8744 | 0.8881 |
| 256 | 0.8806 | 0.9002 |
| 512 | 0.8760 | 0.8954 |

#### 3) Comparison with unsupervised clustering

To highlight the advantages of the weakly supervised vector quantization module, we conduct a comparison experiment by replacing it with K-Means, which is a type of unsupervised clustering. In this setup, each embedding vector is replaced by its corresponding cluster centroid. As an unsupervised algorithm, K-Means is not task-specific and merely converts general embeddings from continuous to discrete. Table V shows the results of K-Means on Camelyon16 under different value of K. It yields AUC that is about 20% lower and the accuracy is about 15% lower compared to the vector quantization module.

**TABLE V:** KMeans (Unsupervised Clustering) Results on Camelyon16.

| K | Accuracy | AUC |
| --- | --- | --- |
| 2 | 0.6946 | 0.6976 |
| 8 | 0.7054 | 0.7040 |
| 16 | 0.6791 | 0.6438 |
| 32 | 0.7318 | 0.6944 |
| 64 | 0.7333 | 0.6887 |
| 128 | 0.7318 | 0.6982 |

#### 4) Impact of different pseudo-bag size

Table VI shows the slide-level performance on Camelyon16 concerning different sizes of pseudo-bags. When the pseudo-bag size is too small, performance decreases rapidly. Since slides in Camelyon16 usually contain only a tiny fraction of tumor tissues, a small pseudo-bag may not include any positive instances. The best performance is achieved when the pseudo-bag size is 256. Performance decreases when the size of the pseudo-bags becomes larger because larger bags contain many simple instances, which are not helpful for the model to identify challenging instances.

**TABLE VI:** Different size of the pseudo bags on Camelyon16.

| size | Accuracy | AUC |
| --- | --- | --- |
| 128 | 0.8031 | 0.8119 |
| 256 | 0.8837 | 0.9248 |
| 512 | 0.8589 | 0.8859 |
| 8192 | 0.8558 | 0.8294 |

## V. Conclusion and Future Works

In this paper, we proposed VQ-MIL, a novel method that integrates weakly supervised vector quantization into the MIL framework for WSI classification. By transforming continuous embeddings into discrete, task-specific codewords, VQ-MIL addresses two challenges in WSI classification, the lack of task-specificity and the high variability associated with continuous embeddings. Our experiments demonstrate that VQ-MIL not only achieves state-of-the-art classification results on the Camelyon16 and TCGA lung cancer dataset but also improves the interpretability by providing clearer instance-level annotations.

Beyond pathology, we believe that the idea of VQ-MIL introduced in our work has broader applications in other weakly supervised tasks where handling large-scale data with limited annotations is common.

In future work, we plan to explore the potential of extending VQ-MIL’s discretization approach to more granular instance-level classification. We also aim to investigate optimizing the codebook design and exploring other vector quantization techniques to further enhance model performance and applicability.

## Acknowledgment

The computing resources in this work were provided by Human Genome Center, the Institute of Medical Science, the University of Tokyo.

We would like to thank Yohei Okubo and Yuxuan Pang for their valuable assistance and contributions to this research.

## Notes

### Competing Interest Statement

The authors have declared no competing interest.

### Summary of Updates

Rephrase the description and add new experiment results.

